# Nelfinavir markedly improves lung pathology in SARS-CoV-2-infected Syrian hamsters despite lack of an antiviral effect

**DOI:** 10.1101/2021.02.01.429108

**Authors:** Caroline S. Foo, Rana Abdelnabi, Suzanne J. F. Kaptein, Xin Zhang, Sebastiaan ter Horst, Raf Mols, Leen Delang, Joana Rocha-Pereira, Lotte Coelmont, Pieter Leyssen, Kai Dallmeier, Valentijn Vergote, Elisabeth Heylen, Laura Vangeel, Arnab K. Chatterjee, Pieter Annaert, Patrick Augustijns, Steven De Jonghe, Dirk Jochmans, Birgit Weynand, Johan Neyts

## Abstract

In response to the ongoing COVID-19 pandemic, repurposing of drugs for the treatment of SARS-CoV-2 infections is being explored. The HIV protease inhibitor Nelfinavir, widely prescribed in combination with other HIV inhibitors, has been shown to inhibit *in vitro* SARS-CoV-2 replication. We here report on the effect of Nelfinavir in the Syrian hamster SARS-CoV-2 infection model. Although treatment of infected hamsters with either 15 or 50 mg/kg BID Nelfinavir [for four consecutive days, initiated on the day of infection] does not reduce viral RNA loads nor infectious virus titres in the lungs compared to the vehicle control, the drug reduced virus-induced lung pathology to nearly the baseline scores of healthy animals. A substantial interstitial infiltration of neutrophils is observed in the lungs of treated (both infected and uninfected) animals. The protective effect of Nelfinavir on SARS-CoV-2-induced lung pathology (at doses that are well tolerated and that result in exposures nearing those observed in HIV-infected patients) may lay the foundation for clinical studies with this widely used drug.

## Introduction

The severe acute respiratory syndrome coronavirus 2 (SARS-CoV-2) resulted in more than 2.0 million deaths within the time-span of a year after its discovery (1). SARS-CoV-2-induced COVID-19 is mainly an inflammatory disease that displays a variety of symptoms from mild (fever, dry cough, and anosmia) to severe (massive cytokine storms and exaggerated immune responses, resulting in severe complications such as acute respiratory distress syndrome (ARDS)) (2). Given the lack of specific therapeutic/prophylactic coronavirus-specific drugs and the current urgency of the situation, repurposing of clinically approved/evaluated drugs is being adopted for rapid identification of treatment for COVID-19 (3).

Nelfinavir mesylate (Viracept), an FDA-approved HIV-protease inhibitor (PI) (4), prevents the cleavage of HIV Gag and Gag-Pol polyproteins and results in immature and non-infectious virus particles (5). Nelfinavir has demonstrated *in vitro* inhibitory activity against SARS-CoV-2 (6) and inhibition of SARS-CoV-2 Spike glycoprotein-mediated cell fusion (7). *In silico* docking studies predict binding of Nelfinavir at the catalytic site of the SARS-CoV-2 main protease, M^pro^ (8).

We here wanted to assess whether Nelfinavir results in antiviral effect against SARS-CoV-2 in Syrian hamsters, which are highly susceptible to infection with the virus and develop a COVID-like lung pathology (9), and therefore is well-suited to assess the impact of antiviral drugs (such as Favipiravir and Molnupiravir (EIDD-2801)) (9, 10).

## Results

Six to eight weeks female Syrian Golden hamsters were treated with Nelfinavir (15 mg/kg/dose or 50 mg/kg/dose) or the vehicle control by intraperitoneal injection (IP) within one hour before intranasal infection with 50 μl containing 2×10^6^ TCID_50_ of SARS-CoV-2 [BetaCov/Belgium/GHB-03021/2020 (EPI ISL 109 72 407976|2020-02-03)] (Fig. 1A). Treatment was administered twice daily for four consecutive days. At 4 d.p.i, lungs were collected for viral RNA load, infectious virus titres quantification, and lung histopathology as previously described (9). Treatment with either 15 mg/kg BID or 50 mg/kg BID Nelfinavir resulted in comparable levels of lung viral RNA load and infectious virus titres as the vehicle control group (Fig. 1B, C), indicating a lack of *in vivo* antiviral efficacy of Nelfinavir in this model. Hamsters in the control and 15 mg/kg BID Nelfinavir-treated groups maintained relatively stable weights throughout the study, whereas an average weight loss of about 7% was observed in the 50 mg/kg BID Nelfinavir-treated group compared to the vehicle group at 0 dpi (Fig. 1D). This weight loss can be attributed to the combination of SARS-CoV-2 infection and the administration of 50 mg/kg BID Nelfinavir, as this dose alone did not affect the weight of two uninfected hamsters (data not shown). No other obvious adverse effects were observed in either of the treatment groups. Remarkably, a significant three-fold reduction in the overall histological disease score was observed in the lungs of infected animals treated with 50 mg/kg BID Nelfinavir as compared to the placebo control. In fact, the histopathology score in the treated group was comparable to the baseline score in untreated, non-infected hamsters (median score of 1.25) (Fig. 2A). Yet, despite this remarkable beneficial effect on lung pathology, an unusual interstitial infiltration of neutrophils was noted in the treated animals (Fig. 2A, 2B). Two uninfected hamsters treated with 50 mg/kg Nelfinavir had histopathology scores at the baseline of 1 but presented as well with neutrophil infiltration (data not shown), confirming that these intriguing observations are indeed attributed to Nelfinavir treatment.

**Fig. 1.**
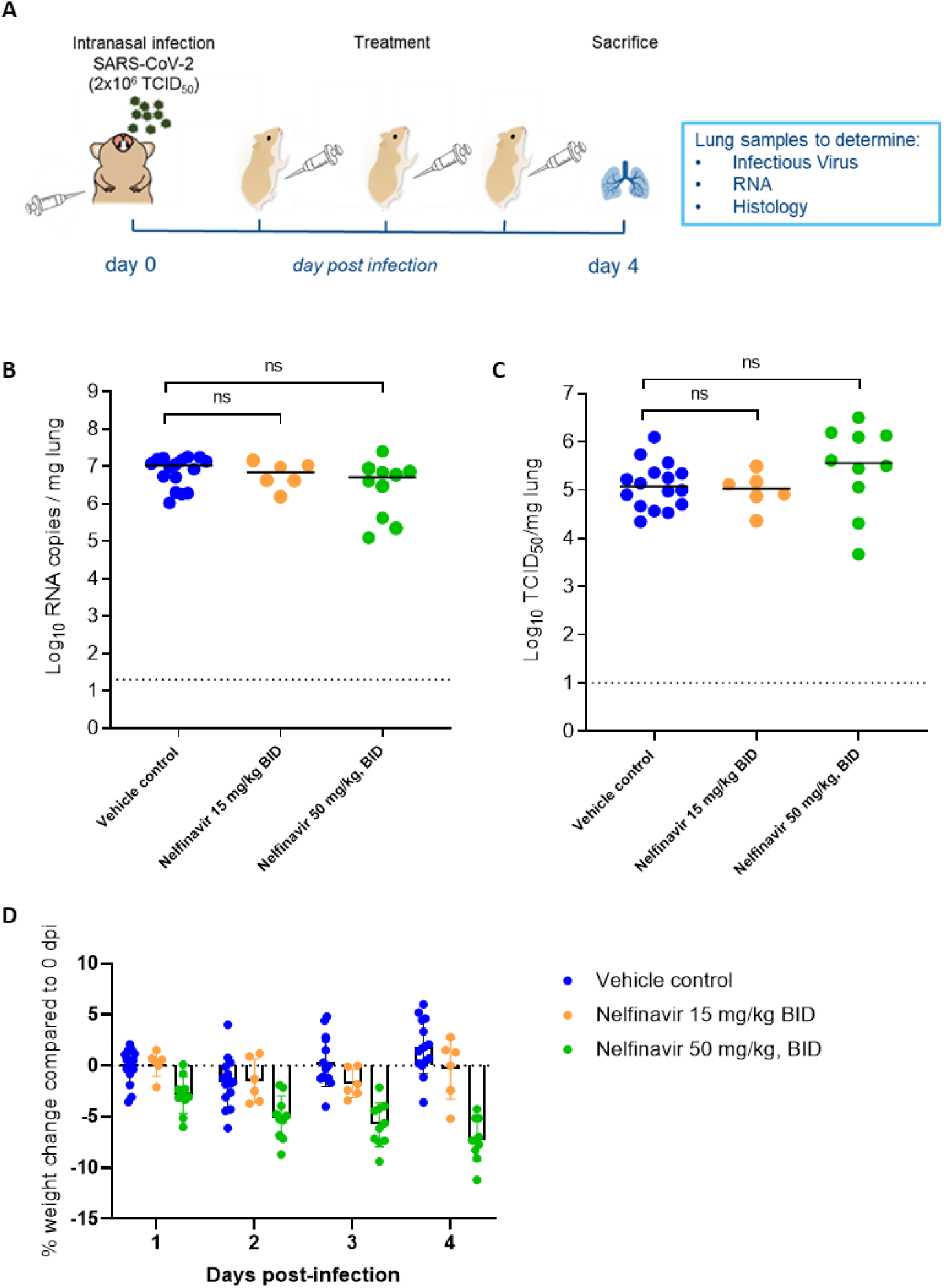
Nelfinavir does not reduce viral replication in SARS-CoV-2 infected hamsters. **(A)** Set-up of the study. **(B)** Viral RNA levels in the lungs of hamsters treated with vehicle (control), 15 mg/kg Nelfinavir BID, or 50 mg/kg Nelfinavir BID at 4 dpi expressed as log_10_ SARS-CoV-2 RNA copies per mg lung tissue. Individual data and median values are presented. **(C)** Infectious virus titers in the lungs of hamsters treated with vehicle (control), 15 mg/kg Nelfinavir BID, or 50 mg/kg Nelfinavir BID at 4 dpi expressed as log_10_ TCID_50_ per mg lung tissue. Individual data and median values are presented. **(D)** Daily weight change in percentage, normalized to the body weight at the time of infection. Bars represent average ± SD. All data (panels B, C, D) are from one (for 15 mg/kg dose) to two independent experiments with a total of 16, 6, and 10 animals for the placebo, 15 mg/kg, and 50 mg/kg conditions respectively. Data were analyzed with the Mann-Whitney U test. ns, non-significant.

**Fig. 2.**
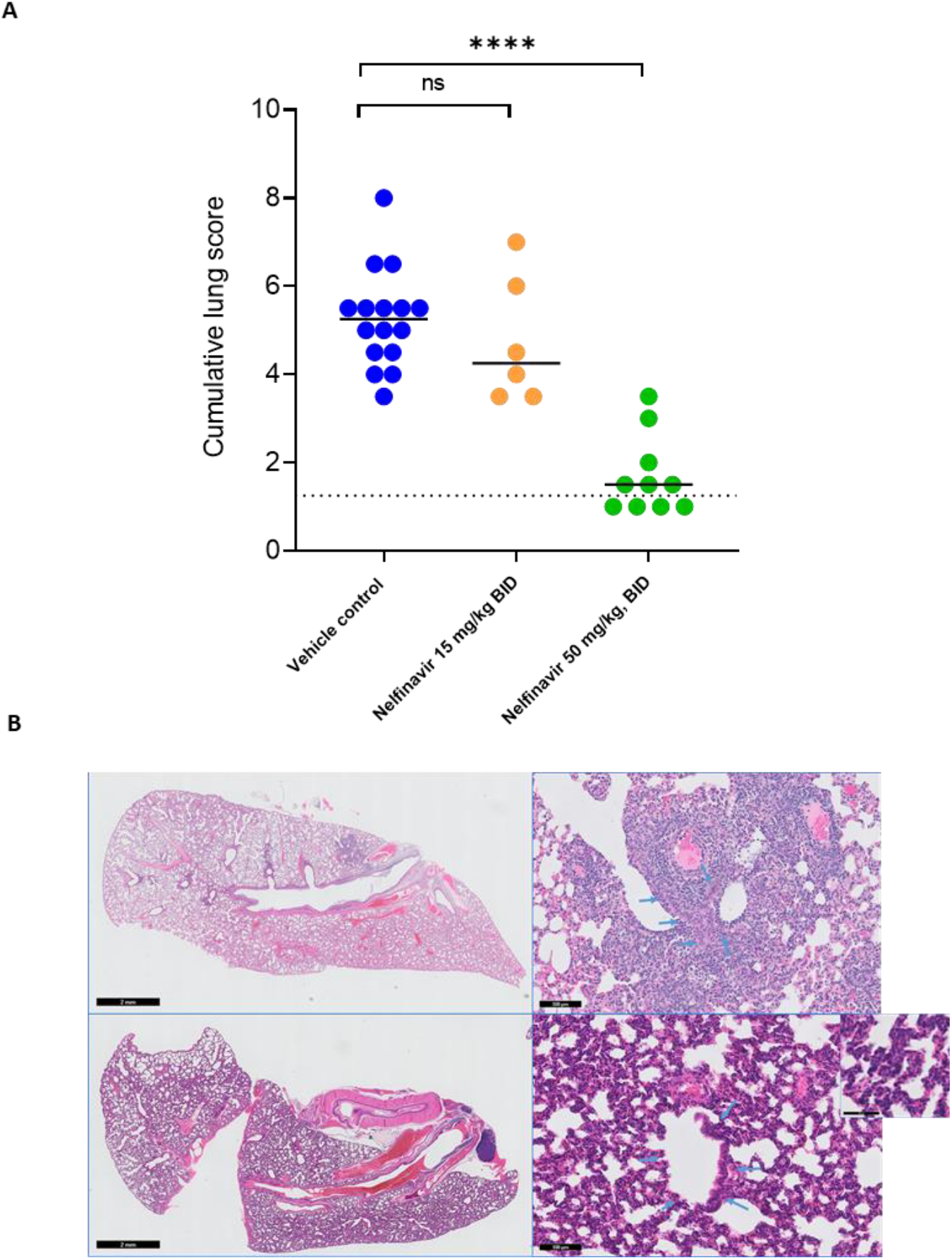
Nelfinavir markedly improves lung pathology of SARS-CoV-2-infected Syrian hamsters despite interstitial neutrophil infiltration. (A) Cumulative severity score from H&E-stained slides of lungs from vehicle control and Nelfinavir (50 mg/kg BID)-treated SARS-CoV-2-infected hamsters at 4 dpi. Individual datapoints and lines indicating median values are presented. The dotted line represents the median lung score in healthy untreated non-infected animals. Data were analyzed using the Mann-Whitney U test. ****P<0.0001. (B) Representative H&E-stained slides of lungs from vehicle control (top panels) and Nelfinavir-treated (bottom panels) SARS-CoV-2-infected hamsters at 4 dpi. Top, left: peri-bronchial inflammation and bronchopneumonia in vehicle-treated hamsters. Top, right: higher magnification of bronchopneumonia centred on a bronchiol (blue arrows) in vehicle-treated hamsters. Bottom, left: dark blue-stained interstitium and bronchial structures appearing normal without inflammation in Nelfinavir-treated hamsters. Bottom, right: severe interstitial inflammation with mostly PMN and congestion. Blue arrows indicate bronchiol (Insert at higher magnification).

At the time of sacrifice (16 h after the last treatment), an average Nelfinavir plasma concentration of 174 nM was measured in 50 mg/kg BID-treated hamsters.

Taken together, although Nelfinavir does not reduce viral load in the lungs of SARS-CoV-2-infected hamsters, it markedly improves lung SARS-CoV2-induced pathology despite a massive infiltration of neutrophils.

## Discussion

The lack of an antiviral activity of Nelfinavir against SARS-CoV2 in hamsters as observed here is consistent with preliminary unpublished data in which Nelfinavir [at 250 mg/kg/day BID orally and boosted with Ritonavir] had no effect on viral lung titers in hamsters. However, no histopathological analysis was reported (Nougairede et al., 2020 – interim report).

HIV PIs exert multiple effects on various cellular processes, such as reducing apoptosis of T cells and neutrophils (11). In particular, Nelfinavir was reported to markedly inhibit apoptosis of neutrophils and neutrophil function through inhibition of calpain (12). These findings, made in the context of HIV, are consistent with our observations in Nelfinavir-treated SARS-CoV-2-infected hamsters. As part of Highly Active Antiretroviral Therapy (HAART) treatment, Nelfinavir is dosed at 1250 mg BID or 750 mg TID, resulting in a C_trough_ of about 370 nM measured after 12 h in patients (13). The Nelfinavir C_trough_ plasma concentrations that are obtained here in hamsters with a 50 mg/kg dose at 16 hr post dosing (on average 174 nM) are therefore close to or in a clinically relevant range.

Neutrophils extracellular traps (NETs) and matrix metalloproteinase-9 (MMP-9), which is a neutrophil-released enzyme, are significantly increased in COVID-19 patients (14). Moreover, the activation of NETs in COVID-19 patients was reported to correlate with disease severity and to be associated with thrombo-inflammation events (14). Consequently, the inhibition of neutrophil function by Nelfinavir and hence downstream immune signalling cascades may contribute to the observed reduction of SARS-CoV-2-induced pathology as observed in hamsters. Furthermore, the reduction of neutrophil apoptosis by the compound may account for the unusually large neutrophil infiltration observed in the lung interstitium in treated animals.

The protective effect of Nelfinavir on SARS-CoV2-induced lung pathology observed here may lay the foundation for clinical studies to explore whether a similar protective activity may be achieved in COVID patients. In the case that the drug would also protect patients against lung damage, combination therapy with directly-acting antivirals (such as Favipiravir and Molnupiravir) against SARS-CoV-2 should also be considered.

## Materials and Methods (Supporting Information)

### SARS-CoV-2 strain

The SARS-CoV-2 strain used in this study, BetaCov/Belgium/GHB-03021/2020 (EPI ISL 109 407976|2020-02-03), was recovered from a nasopharyngeal swab taken from an RT-qPCR confirmed asymptomatic patient who returned from Wuhan, China in the beginning of February 2020. A close relation with the prototypic Wuhan-Hu-1 2019-nCoV (GenBank accession 112 number MN908947.3) strain was confirmed by phylogenetic analysis. Infectious virus was isolated by serial passaging on HuH7 and Vero E6 cells (9); passage 6 virus was used for the study described here. The titer of the virus stock was determined by end-point dilution on Vero E6 cells by the Reed and Muench method (15). Live virus-related work was conducted in the high-containment A3 and BSL3+ facilities of the KU Leuven Rega Institute (3CAPS) under licenses AMV 30112018 SBB 219 2018 0892 and AMV 23102017 SBB 219 20170589 according to institutional guidelines.

### Cells

Vero E6 cells (African green monkey kidney, ATCC CRL-1586) were cultured in minimal essential medium (Gibco) supplemented with 10% fetal bovine serum (Integro), 1% L- glutamine (Gibco) and 1% bicarbonate (Gibco). End-point titrations were performed with medium containing 2% fetal bovine serum instead of 10%.

### SARS-CoV-2 infection model in hamsters

The hamster infection model of SARS-CoV-2 has been described before (9). In brief, wild-type Syrian Golden hamsters (*Mesocricetus auratus*) were purchased from Janvier Laboratories and were housed per two in ventilated isolator cages (IsoCage N Biocontainment System, Tecniplast) with ad libitum access to food and water and cage enrichment (wood block). The animals were acclimated for 4 days prior to study start. Housing conditions and experimental procedures were approved by the ethics committee of animal experimentation of KU Leuven (license P065-2020). Female hamsters of 6-8 weeks old were anesthetized with ketamine/xylazine/atropine and inoculated intranasally with 50 μL containing 2×10^6^ TCID50 SARS-CoV-2 (day 0).

### Study design

For D0 treatment, animals were treated twice daily with intraperitoneal injection (i.p.) of 15 mg/kg or 50 mg/kg of Nelfinavir (purchased from MedChem Express, formulated in 5% DMSO-5% PEG-400 and 5% Tween-80 in PBS to a stock of 5 mg/ml and 10 mg/ml respectively), or vehicle control (5% DMSO-5% PEG-400 and 5% Tween-80) just before infection with SARS-CoV-2. Treatments continued until day 3 pi. During this time, hamsters were monitored for appearance, behavior, and weight. At day 4 pi, hamsters were euthanized by i.p. injection of 500 μL Dolethal (200mg/mL sodium pentobarbital, Vétoquinol SA). Lungs were collected and viral RNA and infectious virus were quantified by RT-qPCR and end-point virus titration, respectively.

### SARS-CoV-2 RT-qPCR

Hamster lung tissues were collected after sacrifice and were homogenized using bead disruption (Precellys) in 350 μL RLT buffer (RNeasy Mini kit, Qiagen) and centrifuged (10.000 rpm, 5 min) to pellet the cell debris. RNA was extracted according to the manufacturer’s instructions. Of 50 μL eluate, 4 μL was used as a template in RT-qPCR reactions. RT-qPCR was performed on a LightCycler96 platform (Roche) using the iTaq Universal Probes One-Step RT-qPCR kit (BioRad) with N2 primers and probes targeting the nucleocapsid (9). Standards of SARS-CoV-2 cDNA (IDT) were used to express viral genome copies per mg tissue or per mL serum.

### End-point virus titrations

Lung tissues were homogenized using bead disruption (Precellys) in 350 μL minimal essential medium and centrifuged (10,000 rpm, 5min, 4°C) to pellet the cell debris. To quantify infectious SARS-CoV-2 particles, endpoint titrations were performed on confluent Vero E6 cells in 96-well plates. Viral titers were calculated by the Reed and Muench method (15) using the Lindenbach calculator and were expressed as 50% tissue culture infectious dose (TCID_50_) per mg tissue.

### Histology

For histological examination, the lungs were fixed overnight in 4% formaldehyde and embedded in paraffin. Tissue sections (5 μm) were analyzed after staining with hematoxylin and eosin and scored blindly for lung damage by an expert pathologist. The scored parameters, to which a cumulative score of 1 to 3 was attributed, were the following: congestion, intra-alveolar hemorrhagic, apoptotic bodies in bronchus wall, necrotizing bronchiolitis, perivascular edema, bronchopneumonia, perivascular inflammation, peribronchial inflammation and vasculitis.

### Statistics

GraphPad Prism (GraphPad Software, Inc.) was used to perform statistical analysis. Statistical significance was determined using the non-parametric Mann Whitney U-test. P-values of ≤0.05 were considered significant.

## Acknowledgments

We thank Carolien De Keyzer, Lindsey Bervoets, Thibault Francken, Elke Maas, Jasper Rymenants, Birgit Voeten, Dagmar Buyst, Niels Cremers, Bo Corbeels and Kathleen Van den Eynde for excellent technical assistance, Piet Maes for kindly providing the SARS-CoV-2 strain used in this study, Jef Arnout and Annelies Sterckx, and the Animalia and Biosafety Departments of KU Leuven for facilitating the animal studies.

